# How People Initiate Energy Optimization and Converge on Their Optimal Gaits

**DOI:** 10.1101/503433

**Authors:** Jessica C. Selinger, Jeremy D. Wong, Surabhi N. Simha, J. Maxwell Donelan

## Abstract

A central principle in motor control is that the coordination strategies learned by our nervous system are often optimal. Here we combined human experiments with computational reinforcement learning models to study how the nervous system navigates possible movements to arrive at an optimal coordination. Our experiments used robotic exoskeletons to reshape the relationship between how participants walk and how much energy they consume. We found that while some participants used their relatively high natural gait variability to explore the new energetic landscape and spontaneously initiate energy optimization, most participants preferred to exploit their originally preferred, but now suboptimal, gait. We could nevertheless reliably initiate optimization in these exploiters by providing them with the experience of lower cost gaits suggesting that the nervous system benefits from cues about the relevant dimensions along which to re-optimize its coordination. Once optimization was initiated, we found that the nervous system employed a local search process to converge on the new optimum gait over tens of seconds. Once optimization was completed, the nervous system learned to predict this new optimal gait and rapidly returned to it within a few steps if perturbed away. We model this optimization process as reinforcement learning and find behavior that closely matches these experimental observations. We conclude that the nervous system optimizes for energy using a prediction of the optimal gait, and then refines this prediction with the cost of each new walking step.

## INTRODUCTION

People often learn optimal coordination strategies. That is, the nervous system tracks a cost for movement and it adapts its coordination to minimize this cost. This optimization principle underlies theories on the control of reaching, grasping, gaze and gait, although the nervous system may seek to minimize different costs for each of these tasks [1-9]. It has provided insight into healthy and pathological behaviour [10-12], as well as the functions of different brain areas [13]. While there is a growing body of evidence that preferred behaviour in these various tasks indeed optimizes sensible cost functions [1-9,14,15], how the nervous system performs this optimization is largely unknown [2,16].

The optimization of movement is a challenge for the nervous system. To perform a movement, the nervous system has thousands of motor units at its disposal, and it can finely vary each motor unit’s activity many times per second. This flexibility results in a combinatorially huge number of candidate control strategies for performing most movements—far too many for the nervous system to simply try each one to evaluate its cost [17,18]. The nervous system must instead efficiently search through its options to seek optimal solutions within usefully short periods of time. A second consequence of the large number of control strategies available to the nervous system is that it can never know whether it has truly converged to the best of all possible options. But if it is indeed at an optimum, continuously searching for better options will itself be sub-optimal because all other executed coordination patterns will be more costly [19]. Thus, the nervous system must determine when to initiate optimization and explore new coordination patterns, and when to exploit previously learned strategies [20-22].

Here we use human walking to understand how the nervous system initiates and performs the optimization of its motor control strategies. Human walking is a system well suited for studying these questions because the primary contributor to the nervous system’s cost—metabolic energy expenditure—is both well established and measurable in this task. Decades of experiments that used respiratory gas analysis have established that our preferred gait parameters—from walking speed [23-26] to step frequency [24,27-29] and step width [30]—minimize energetic cost. While some optimal motor control strategies may be established over relatively long periods of time, we recently discovered that the nervous system can re-optimize aspects of gait within minutes [31]. This is a second reason why human walking is well suited for studying optimization—we can observe energy optimization within a lab setting and within a reasonably short period of time. Studying optimization in tasks such as reaching or saccades is less straightforward as the relevant cost to the nervous system appears to include a term not only for task effort, but also for the task error, and with some unknown weighting between these two contributors [2,5,32,33]. Furthermore, motor learning in these tasks appears to, at least initially, prioritize reducing error over optimizing cost, requiring creative experiments to decouple error-based learning from reward-based learning [10,34].

To study how the nervous system performs energy optimization in human walking, we leveraged our previously-developed experimental design within which people reliably optimize their gait to minimize energetic cost [31]. In brief, we used lightweight robotic exoskeletons capable of applying resistive torques at the knee joints during walking (**Fig 1**). The exoskeleton controller applies a resistance, and therefore an added energetic penalty, that is minimal at low step frequencies and increases as step frequency increases. Our past experiments demonstrated that this control function reshapes the relationship between step frequency and energetic cost—which we term the *cost landscape*—creating a positively sloped energetic gradient at subjects’ initial preferred step frequency, and an energetic minimum at a lower step frequency. When given sufficient experience with the new cost landscape, subjects in our past experiments learned to decrease their step frequency to converge on the new energetic minimum. We use the term optimization to refer to the process of adapting coordination towards new patterns that minimize cost. This might alternatively be called reward-based adaptation [10,34]. We also distinguish between optimization and prediction, where the former is the process of trying new coordination patterns as the nervous system converges towards the minimum cost, and the latter is the nervous system storing and recalling previously experienced coordination patterns [35,36]. For our purposes we consider prediction, because it involves the storing of a coordination pattern, as commensurate with learning.

**Fig 1.**
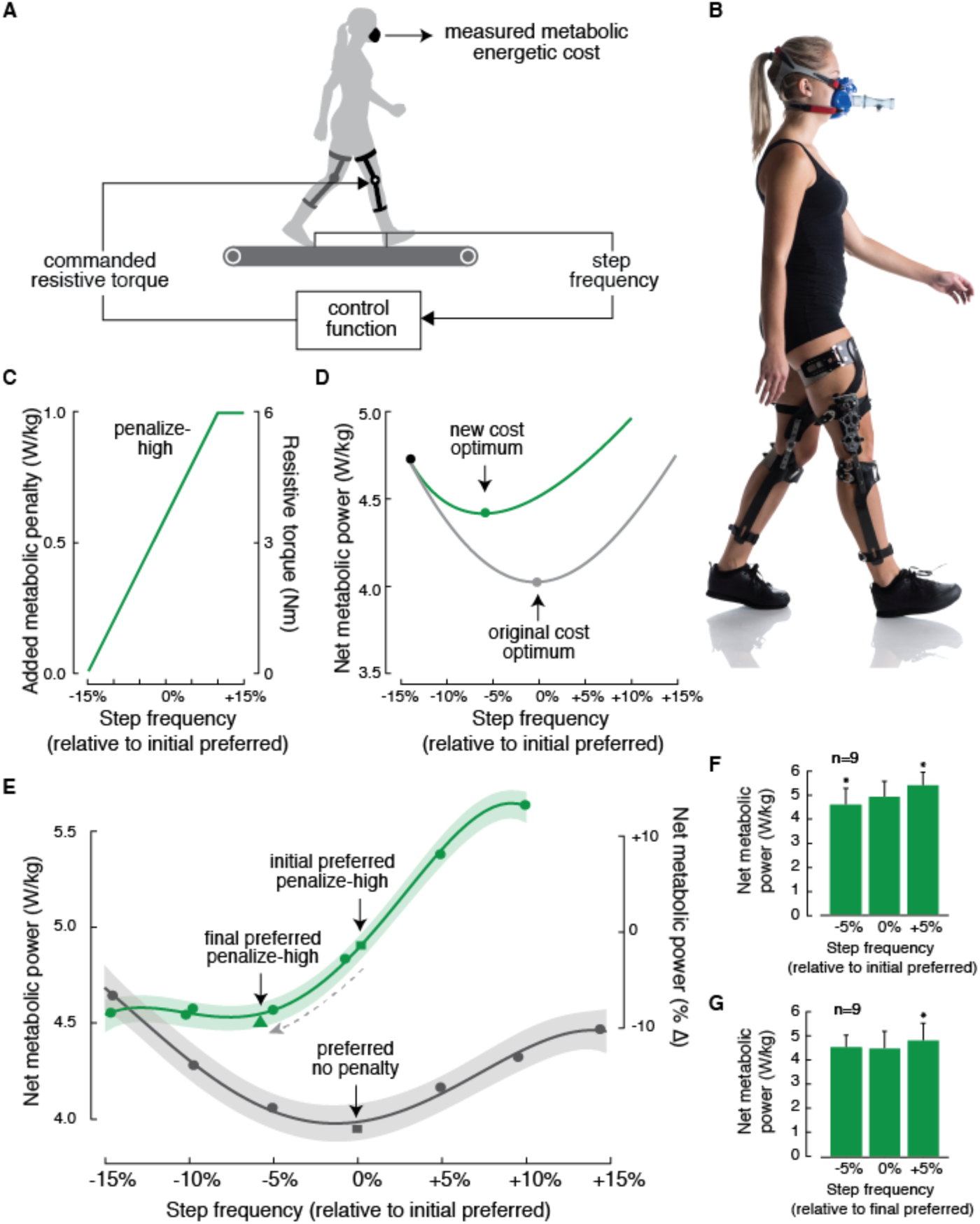
Experimental design. (**A-B**) By controlling a motor attached to the gear train of our exoskeletons, we can apply a resistance to the limb that is proportional to the subject’s step frequency. (**C**) Design of the penalize-high (green) control function. (**D**) Schematic energetic cost landscapes. Adding the energetic cost of the penalize-high control function to the original cost landscape (grey) produces a new cost landscape with the optimum shifted to lower step frequencies (green curve). (**E**) Measured energetic cost landscapes, reproduced from Selinger et al. (2015), for the penalize-high (green) control function and controller off condition (grey). The lines are 4^th^ order polynomial fits, and the shading their 95% confidence intervals, shown only for illustrative purposes. The dashed grey arrow illustrates the direction of adaptation from initial preferred (green square) to final preferred step frequencies (green triangle). On average subjects decreased their step frequency by approximately 6% to converge on the energetic minima and reduce cost by 8%. (**F**) The penalize-high control function creates a positively sloped energetic gradient about the subjects’ initial preferred step frequency. (**G**) Subjects adapted their step frequency to converge on the energetic minima. Error bars represent 1 standard deviation. Asterisks indicate statistically significant differences in energetic cost when compared to the cost at the initial or final preferred step frequency (0%).

Although our prior experiment was the first to provide direct experimental evidence for continuous energy optimization [31], it did not allow us to decipher what experience with a novel cost landscape is critical for optimization to be initiated and what process is used to converge on optima. To understand these mechanisms, here we used a series of experiments that controlled the type of initial experience subjects received with a new energetic cost to determine what gait experience was sufficient for the nervous system to stop exploiting a previously optimal solution and initiate a new optimization. Once the nervous system initiated optimization, we studied how it explored new gaits, in order to understand the nervous system’s algorithms for converging on new energetic optima. Using the results of our experiments that examined both the initiation and process of optimization, we then developed computational models based on reinforcement learning that explain how the nervous system may optimize energy during walking.

## RESULTS

### High natural gait variability may spontaneously initiate optimization

We first sought to determine whether the nervous system could spontaneously initiate optimization and converge on new energetic optima. All subjects first walked for 12 minutes while wearing the exoskeletons, but with the controller turned off (**Fig 2A**, Baseline). This meant that while there was some small inertial and frictional torques from the exoskeleton, there was no additional resistive torque added by the robotic motor [31]. All walking took place on an instrumented treadmill at 1.25m/s and we measured step frequency from treadmill foot contact events [37]. All subjects appeared to settle into a steady state step frequency within 9 minutes and we used the final 3 minutes of walking data to determine subjects’ ‘initial preferred step frequency’. On average, subjects walked at 1.8 ± 0.1 Hz (mean ± SD). To guard against the possibility that in future trials subjects could be unaware they are able to alter, or fearful to alter, their step frequency when walking on a treadmill at a constrained speed, we next habituated subjects to walking at a range of step frequencies (**Fig 2B**, Habituation). During this habituation, the controller remained off; therefore, subjects did not gain experience with the new cost landscape. We then turned the controller on, resulting in an applied resistance that was dependent on step frequency, and the subjects walked at a self-selected step frequency for an additional 12 minutes (**Fig 2C**, First experience). During this first experience period, 6 of the 36 subjects displayed gradual adaptations in gait and converged to lower, less costly, step frequencies consistent with the energetic optima (**Fig 3A-B**). These subjects, whom we refer to as ‘spontaneous initiators’, had to meet two criteria. First, during the final 3-minutes of the first experience period their average step frequency was required to fall below 3 SD in steady state variability, determined from the final 3-minutes of the baseline period. For most subjects, this equates to a minimum decrease in step frequency of approximately 5%. Second, the change in step frequency could not be an immediate and sustained mechanical response to the exoskeleton turning on. Subjects’ final step frequency had to be significantly lower than the step frequency measured in the 10th to 40th second after the exoskeleton turned on (one-tailed t-test, p<0.05). See Supporting Information **Fig S1** for discrimination plot and additional discussion of these criteria. On average, the spontaneous initiators converged toward the optima with an average time constant of 65.7 seconds (exponential fit 95% CI [60.5, 70.8]), or about 120 steps. As determined by our criteria, these *spontaneous initiators* settled on a step frequency that is indistinguishable from the location of the expected optima from our previous experiments [31] (one sample t-test, t(5)=-0.46, *p* = 0.66), while the other subjects, or *non-spontaneous initiators*, remained at their initial preferred step frequency (0.8 ± 2.7 %, **Fig 3D**). We hypothesized that high natural gait variability, which results in a more expansive and therefore more clear sampling of the new cost landscape, would be a predicator of spontaneous initiation. To test this, we analyzed individual subjects’ step-to-step variability prior to the controller even being turned on and found that spontaneous initiators displayed higher variability in step frequency than non-spontaneous initiators (1.5 ± 0.3 % and 1.1 ± 0.3 %, respectively, two sample t-test, t(34)=6.06, *p* = 1.8×10^-2^, **Fig 3C**). This finding that spontaneous initiation was correlated with higher variability, even before the adaptation itself, was in no way predetermined by our criteria. As a second test of the role of step frequency variability in promoting spontaneous initiation, we regressed the amount of adaptation an individual exhibited during the First Experience period against their step frequency variability from the Baseline period (final 3-minutes), for all 36 subjects, and found a weak but significant correlation (R^2^=0.22, *p* =4.0×10^-3^). We expect other factors that vary across individuals, such as the gradient of their cost landscape and their levels of sensory and motor noise, to additionally effect the saliency of the cost landscape, and in turn the likelihood of spontaneous initiation.

**Fig 2.**
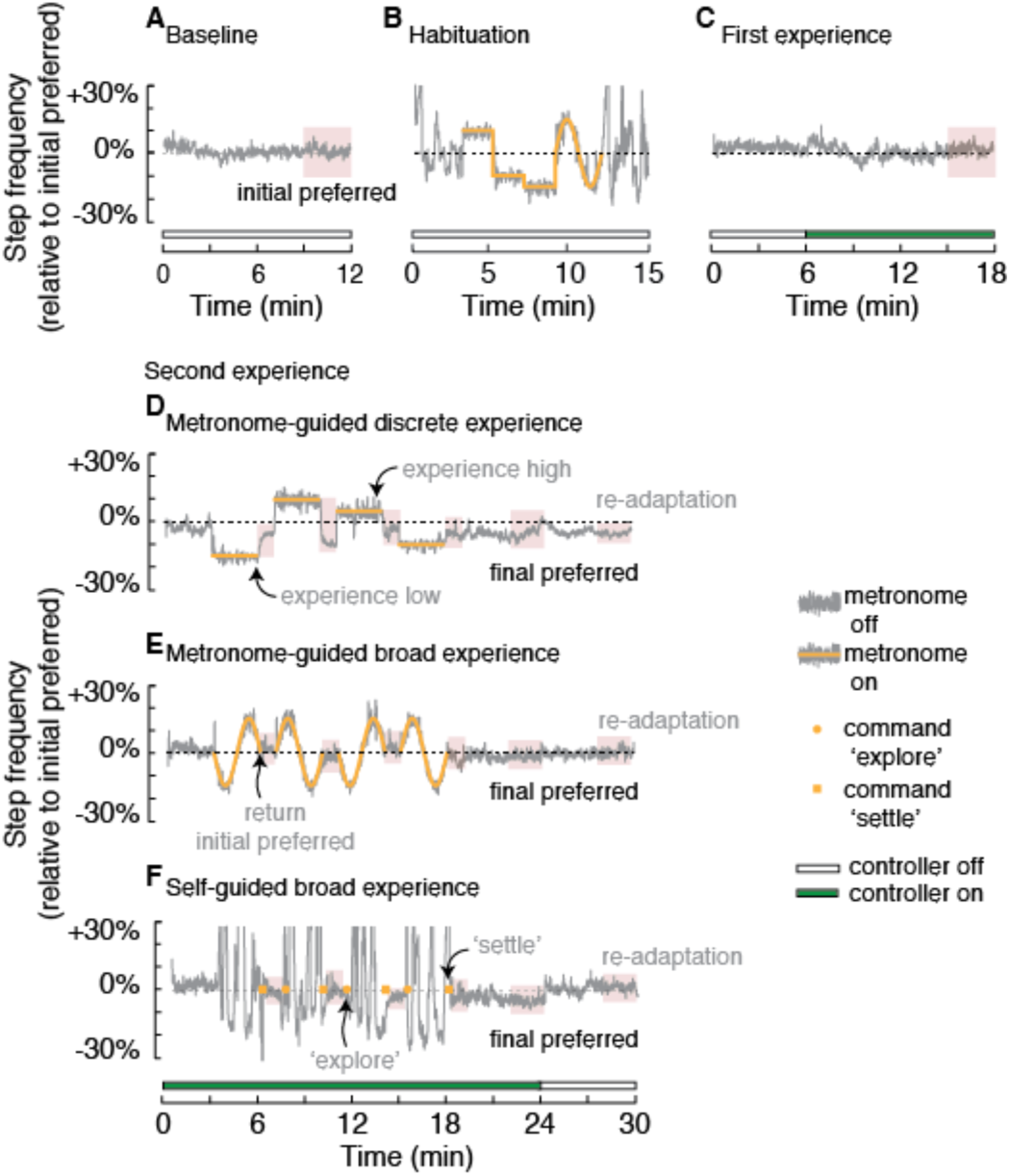
Experimental protocol. Each subject completed four testing periods. The first three, Baseline (**A**), Habituation (**B**), and First Experience (**C**), were the same for all subjects. For the Second Experience period, subjects were assigned to either the metronome-guided perturbations to discrete cost points (**D**), metronome-guided broad experience with the cost landscape (**E**), or self-guided exploratory experience of the cost landscape (**F**). Rest periods of 5-10 minutes were provided between each testing period. For all periods, regions of red shading illustrate the time windows during which we assessed steady-state step frequencies. Data shown in **A-C** and **E** are from one representative subject, while data in **D** and **F** are from two other representative subjects.

**Fig 3.**
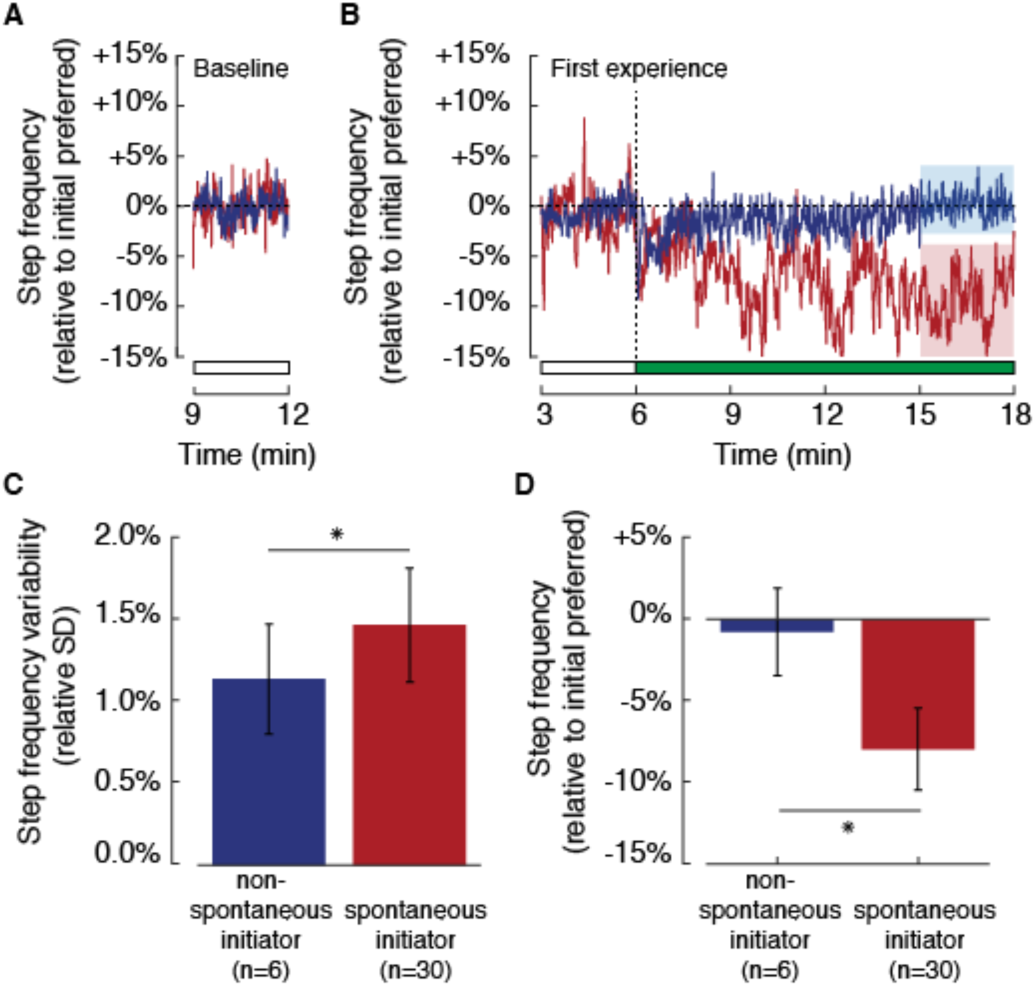
Non-spontaneous and spontaneous initiators. **A**) Self-selected step frequency during the final 3 minutes of the Baseline testing period for representative non-spontaneous initiator and spontaneous initiator (blue and red, respectively). (**B**) Step frequency data during the First Experience period for the same two representative subjects. The horizontal bar indicates when the controller is turned on (green fill) and off (white fill). (**C**) Across all subjects, spontaneous initiators displayed greater average step frequency variability than non-spontaneous initiators during the Baseline testing period. (**D**) By the final 3 minutes of the First Experience period, spontaneous initiators appeared to adapt their step frequency to converge on the energetic minima, while non-spontaneous initiators did not. Error bars represent 1 standard deviation. Asterisks indicate statistically significant differences for t-tests.

### Experience with lower cost gaits can initiate optimization

We next sought to determine how optimization could be initiated in the non-spontaneous initiators. The non-spontaneous initiators were assigned to one of three experiments (**Table S1**) in which a second experience period included either metronome-guided experience with discrete cost points on a new cost landscape (**Fig 2D**), metronome-guided experience with many costs along a new cost landscape (**Fig 2E**), or self-guided experience with many costs along a new cost landscape (**Fig 2F**). To gain insight into the progress of optimization during this period, 1-minute probes of subjects’ self-selected step frequency occurred at the 6^th^, 10^th^, and 14^th^ minute, along with a final 6-minute probe at the 18^th^ minute. We found that if, just prior to the first probe, subjects were walking at low step frequencies, and thus experienced lower energetic costs, they appeared to initiate optimization and adapt toward the new optima (**Fig 4**). Yet, if they were walking at high step frequencies, and thus experienced higher energetic cost, they rapidly returned to the initial preferred step frequency (**Fig 4**). This finding was consistent regardless of whether the experience was self-guided (t-test, t(7)=-2.25, *p* = 0.03) or metronome-guided (t-test, t(7)=-2.33, *p* = 0.03). Moreover, if immediately before the probe subjects were returned to the initial preferred step frequency, as was the case with the metronome-guided experience of many cost points, they showed no adaptation (**Fig 4;** t-test, t(7)=0.12, *p* = 0.55). This was despite them having broad experience with the cost landscape. It appears that providing subjects with experience at a low-cost gait and then allowing them to self-select their gait following these new initial conditions is sufficient for initiating optimizing, while expansive experience with the landscape is not. Importantly, the energy cost at the low-cost gait is lower relative to the energy cost at the initially preferred step frequency under the new cost landscape, but not the original cost landscape (**Fig 1E**) indicating that the nervous system is updating its expectation of the energetic consequences of its gaits.

**Fig 4.**
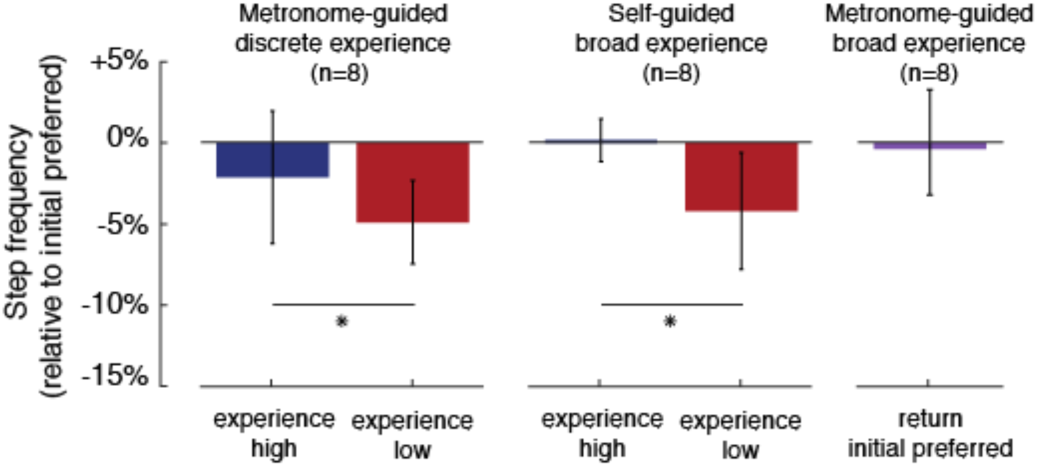
Effect of experience direction on initiation of optimization. For each subject, we averaged data from the final 30 seconds of the first step frequency probe, and then averaged across subjects. Error bars represent 1 standard deviation. Asterisks indicate statistically significant differences for paired t-tests.

### A local search strategy is used to converge on energetically optimal gaits

To investigate the interaction between high and low cost experience, as well as the order of the experience, we next compared the behaviour during the first and final probes following experience with the highest (+10%) and lowest (−15%) step frequencies (**Table S1**). We performed a two-way ANOVA and found that both experience direction (high and low cost) and probe order (first and last) had significant effects (F(1, 17) = 13.25, p = 2.7×10^-3^; F(1, 17) = 4.93, p = 0.04; respectively), as well as their interaction (F(1, 17) = 5.30, p = 0.04). A two-sample t-test revealed that step frequencies were significantly different following the first experience with high and low costs (t(7)=6.1, *p* = 4.8×10^-4^). Following the first experience with high step frequencies subjects appeared to use prediction to rapidly move away from this high cost step frequency (**Fig 5A, Fig S2A**). But, their prediction was erroneous—having not yet experienced lower costs gaits, they returned to their initial preferred step frequency (**Fig 5B, Fig S2A**). They did so with an average time constant of 2.0 seconds (exponential fit 95% CI [1.5 2.5]) or about 4 steps. Following the first experience with low step frequencies subjects more slowly descended the cost gradient, with an average time constant of 10.8 seconds (exponential fit 95% CI [9.2 12.5]), about 20 steps, and eventually converged on the new optima (**Fig 5A**). Because this was the first probe, all of which followed experience at −15% step frequency, these subjects had no prior explicit experience with the new optima yet could converge to it (**Fig 5B**)—prior explicit experience with the new optima was not necessary for convergence. Subjects’ gradual and sequential convergence to the new optima is consistent with a local search process, and inconsistent with alternative optimization methods such as actively sampling from a broad range of new gaits.

**Fig 5.**
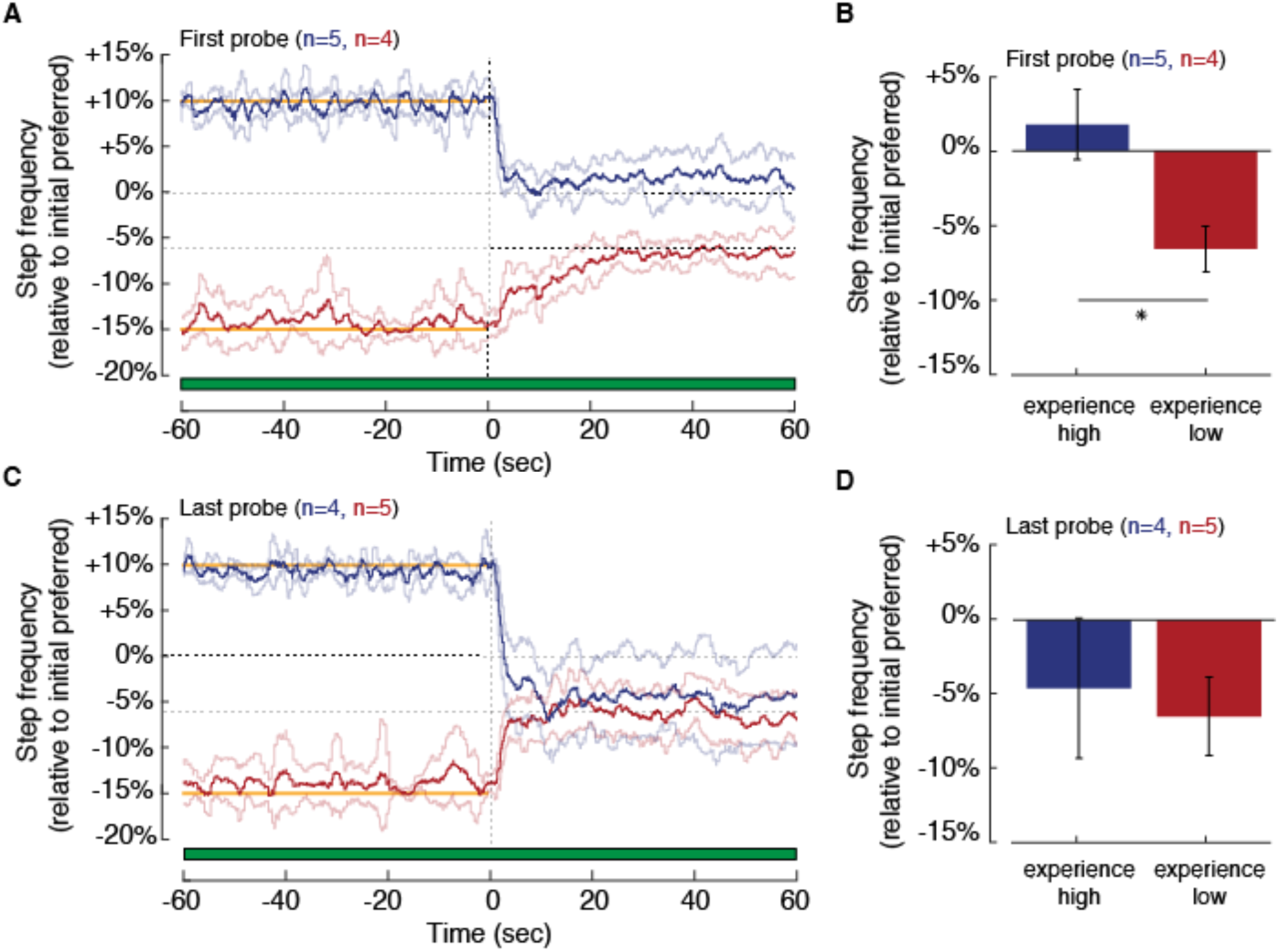
Effect of experience direction during first and last step frequency probe. Step frequency time-series data, averaged across subjects, for the first (**A**) and last (**C**) probes following experience either high (blue) or low (red) step frequencies. The light blue and red lines represent 1standard deviation in step frequency for each time point. The horizontal bars indicate when the controller is turned on (green fill) and off (white fill), and the yellow lines indicate the prescribed metronome frequencies. Steady state step frequencies, averaged across subjects, during the final 30 seconds of the probe for the first (**B**) and last (**D**) perturbations toward either high or low. Error bars represent 1 standard deviation. Asterisks indicate statistically significant differences for t-tests.

### Optimization leads to new predictions of energy optimal gaits

During the last probe, subjects rapidly converge on the new optima, with an average time constant of two to three seconds, regardless of the direction of prior experience (experience high: 2.8 seconds, exponential fit 95% CI [2.3 3.2]; experience low: 2.5 seconds, exponential fit 95% CI [1.9 3.1]; **Fig 5C-D, Fig S2B**). A two-sample t-test revealed that step frequencies were now indistinguishable following the last experience with high and low costs (t(7)=0.77, *p* = 0.47). Following the last experience at low step frequencies, subjects no longer display slow adaptations consistent with optimization, but instead rapidly predict the optimal gait. And, following the last experience at high step frequencies, subjects’ erroneous predictions have been corrected and they rapidly converge to the new cost optimum. This indicates that optimization culminates in the formation of new predictions about optimal movements and the abolishment of old. On average, subjects’ ‘final preferred step frequency’ was −4.8 ± 3.1 %, which was lower than initial preferred (t-test, t(8)=-4.74, *p* = 1.5×10^-3^) and consistent with the expected optima. The high inter-subject variability following the final probe (**Fig 5D**) may in part be due to that fact that each subject will have a different energy optimal step frequencies. When the controller was turned off, returning subjects to the original cost landscape, they slowly unlearned this new prediction. They gradually, with a time constant of 10.5 seconds (exponential fit 95% CI [8.8 12.2]), returned to a step frequency indistinguishable from their initial preferred step frequency when the controller was turned off (−0.8 ± 3.0 %).

### Energy optimization as reinforcement learning

The experimental behaviours we observed, where in a new environment subjects iteratively learn and then rapidly predict the energy optimal gait, resemble the behaviours produced by classic reinforcement learning algorithms [19,38]. As a proof-of-principle for human motor learning, reinforcement learning algorithms have found the optimal control policies for robots and physics-based simulations that walk, reach, and do other movement tasks [39-41]. And, the necessary components to perform reinforcement learning for human movements, including reward prediction and sensory feedback, are present in our nervous systems and well-studied for learning non-motor tasks [42]. Here we test if a simple reinforcement learning model can indeed reproduce the experimental behaviours observed during energy optimization (**Fig 6A**). Reinforcement learning, applied to our context, allows the nervous system to iteratively learn a *value-function* (*Q*) that stores the predicted relationship between step frequency and energetic cost. For each new step, the nervous system selects a step frequency, or *action* (*a*), in accordance with its *policy* (*π*): ‘choose the energy minimal step frequency’. Each time the nervous system executes a frequency, it measures the resulting energetic cost, or *reward* (*r*), and updates its predicted cost for that frequency. However, since the reward can’t be measured perfectly due to *measurement noise* (*n_m_*) nor the action executed perfectly due to *execution noise* (*n_e_*), the nervous system doesn’t simply replace the old predicted value with the new reward. Instead it updates the old value by some fraction of the measured reward, referred to as the *learning rate* (*α*). Our aim here is to demonstrate how a simple reinforcement model of energy optimization can capture the key behavioral features demonstrated by subjects. Our model is not designed to predict or explain individual subjects’ behavior or action histories.

**Fig 6.**
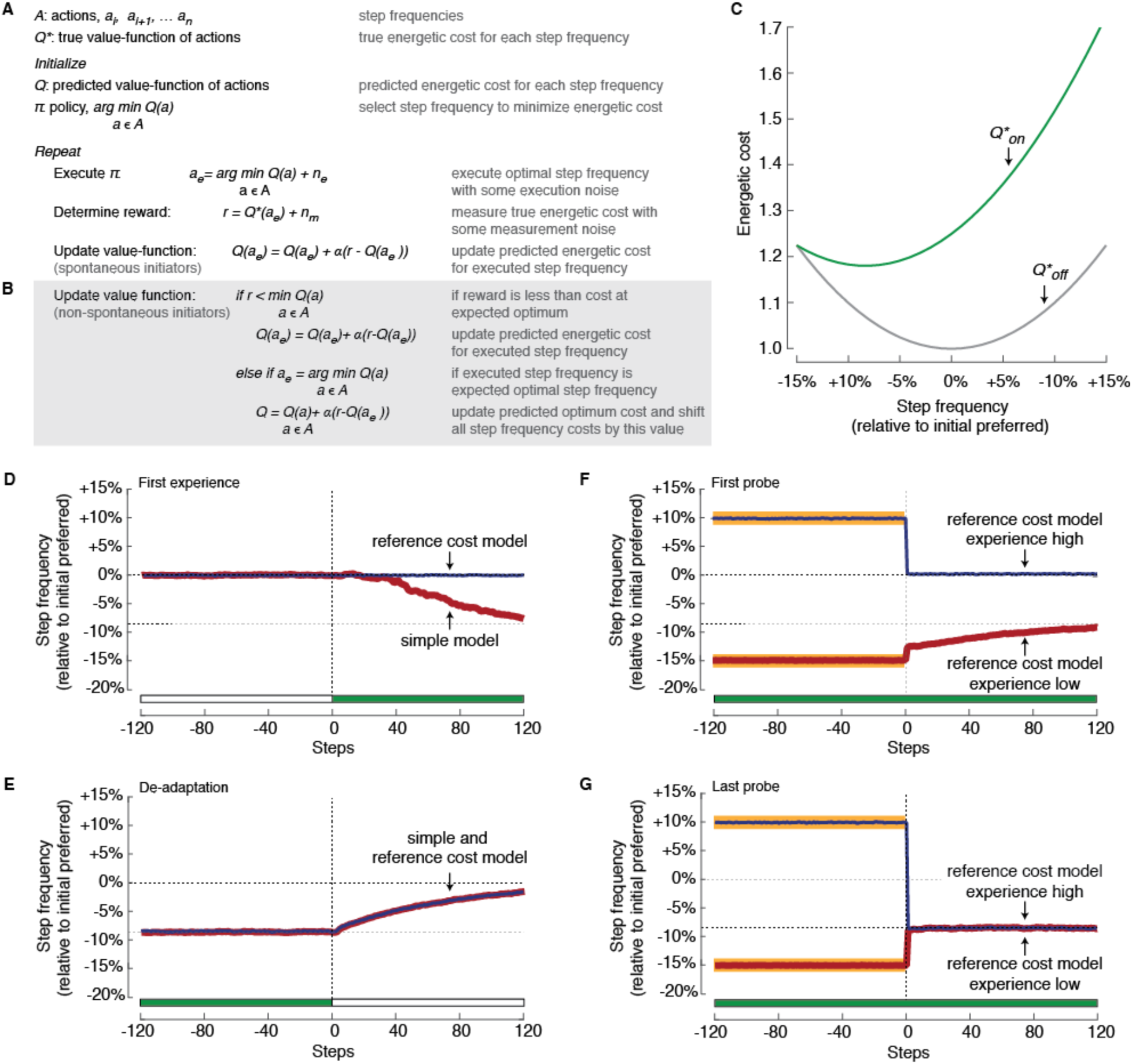
Reinforcement learning model of energy optimization. (**A**) A simple model describing the behavior of spontaneously initiating subjects. (**B**) A more complex logic for updating the value function that prioritizes learning a reference cost and can describe the behavior of non-spontaneously initiating subjects. (**C**) The simulated energetic cost landscapes when the controller is turned off (*Q*_off_*) and on (*Q*_on_*). Behavior of the simple model of spontaneous initiators (red) and reference cost model of non-spontaneous initiators (blue) during the First Experience period when the controller is first turned on (**D**) and the final de-adaptation period when the controller is turned off (**E**). Behavior of the reference cost model of non-spontaneous initiators during the first (**F**) and last (**G**) probes following experience with high (blue) and low (red) step frequencies. The horizontal bars indicate when the controller is turned on (green fill) and off (white fill), and the thick yellow lines indicate the prescribed frequencies prior to the probe.

In this model, we specify the following: i) *Q\** is initially a representation of the dependence of energetic cost on step frequency during natural walking when the controller is turned off—the original cost landscape (*Q*_off_*, **Fig. 6C**); ii) to simulate the controller turning on, we change *Q\** to an accurate representation of the dependence of energetic cost on step frequency under our control function—the new cost landscape (*Q*_on_*, **Fig. 6C**); iii) we represent possible actions as discrete step frequencies; and iv) we set the level of execution noise, measurement noise and learning rate such that the resulting variability in step frequency and rate of convergence to the optimum are comparable to that observed experimentally (See Methods, Model Details). Importantly, the qualitative findings we present below are not particularly sensitive to these specific parameter settings (**Fig S3**).

This very simple model can well describe the behaviour of our spontaneous initiators. We find that over about the same number of steps as our human subjects, the model can converge on new energetically optimal gaits to achieve small cost savings (**Fig 6D**). It also learns to predict the new cost landscape, rapidly returning to new cost optima when perturbed away, just as we have found in our human experiments. When returned to the original and previously familiar cost landscape, it doesn’t instantly remember old optima but instead has to unlearn its new prediction (**Fig 6E**). Notably, our model does not provide insight into individual subject’s behavior, but rather the general behavioural features of energy optimization.

This simple reinforcement learner cannot however explain the behaviour of our non-spontaneous initiators. Unlike the majority of our experimental subjects, the above model will always spontaneously initiate optimization and begin converging on the optimal gait (even if the learning rate is adjusted such that past predictions are much more heavily weighted over new measures, **Fig S3A**). Our experimental findings suggest that non-spontaneous initiators may heavily favour exploitation over exploration [19] until sufficient experience with a low-cost gait signals to the nervous system that the current action is suboptimal (See Methods, Model Details).

One model that can capture this more complicated behaviour of the nervous system is a reinforcement learner that prioritizes the learning of a reference cost [43-45] that equals the cost at the predicted optimum step frequency (**Fig 6B**). This model continuously relearns the value of the reference cost and then shifts the costs associated with all frequencies by this value. The algorithm only recognizes a change to the shape of the cost landscape when it detects a cost saving with respect to this continuously updated reference cost. It then initiates optimization and updates the cost associated with the individual frequencies that it executes, thereby learning the shape of the new cost landscape. Prioritizing the learning of a reference cost, rather than constantly exploring new gaits, is perhaps a better general strategy for cost optimization in real-world conditions. Energetic cost continuously varies as conditions change in the real world, but unlike our experiment, only some conditions may benefit from the adoption of a new gait and exploring gaits away from the optimal gait comes with an energetic penalty. The continuous updating of a reference cost allows the nervous system to detect when there are reliable costs savings to be gained relative to the predicted optimal gait. It also allows the nervous system to compare differences between the two gaits and understand which walking adjustments led to the lower cost [10,46]. This may allow the nervous system to learn the dimension along which exploration should proceed and quickly converge on the new optimal gait [9,10,47].

It is possible that high natural gait variability, as displayed by our spontaneous initiators, is in fact also triggering initiation through the updating of a reference cost because it provides sufficient experience with a low-cost gait. If treated as so, all subjects’ behaviour could be explained by the reference cost model. However, deciphering an exact low-cost experience criterion that fits all subject’s behaviour, is difficult, and perhaps not possible, as it likely varies across subjects and is affected by additional factors such as the gradient of their cost landscape, their levels of sensory and motor noise, and their weighting of newly experienced costs.

This reference cost learning algorithm captures many key behavioral features of our non-spontaneously initiating subjects. First, it does not spontaneously initiate optimization (**Fig 6D**). Second, it only initiates after experience in the new cost landscape with a frequency that has a lower cost than that at the initially preferred frequency. Third, after initiation, the algorithm gradually converges on the new optimum (**Fig 6F**). Finally, much like our original model of spontaneous initiators, after convergence it can leverage prediction to rapidly return to the new optimum after a perturbation (**Fig 6G**) but must slowly unlearn this optimum if returned to the original cost landscape (**Fig 6E**).

Overall, our computational models demonstrate that the nervous system may optimize for energy using algorithms consistent with a reinforcement learning framework—leveraging predictions of the optimal gait, and then refining this prediction with the cost of each new walking step. Our experiments and models allow us to describe three key features of energy optimization during gait. First, initiation of optimization appears to be preferentially triggered by experience with low costs gaits, consistent with a prioritization of the learning of a reference cost [43-45]. Second, during optimization the cost of each new walking step results in an updating of the expected cost landscape. And third, this expected cost landscape allows for rapid prediction, and slow unlearning, of energy optimal gaits.

## DISCUSSION

Here we used energy minimization in human walking to understand how the nervous system initiates and performs the optimization of its motor control strategies. We found that some people tend to explore, through naturally high gait variability, leading them to spontaneously initiate optimization. Others are more likely to exploit their current prediction of the optimal gait and require experience with lower cost gaits to initiate optimization. When optimization was initiated, people gradually adapted their gait, in a manner consistent with a local search strategy, to converge on the new optima. Given more time and experience, this slow optimization was replaced by a new and rapid prediction of the optimal gait. Our reinforcement learning models reproduce these behaviours, suggesting that the nervous system may use similar mechanisms to optimize gait for energy in walking, and perhaps optimize other movements for other cost functions.

Principles of energetic optimality may also determine the nervous system’s balance between exploration and exploitation. Variability can aid with initiation by allowing the nervous system to locally sample a more expansive range of the cost landscape, clarify its estimate of the cost gradient, and identify the most promising dimensions along which to optimize [21,48,49]. This variability may simply be a consequence of noisy sensorimotor control that fortuitously benefits the exploration process, or it may reflect intentional motor exploration by the nervous system [21,48]. Recent work suggesting that humans actively reshape the structure of their motor output variability to elicit faster learning of reaching tasks, is evidence of the latter [48]. Learning rate also affects variability because new cost measurements are imperfect. The higher the learning rate, the greater the influence of the new and noisy cost measurements on the predicted optimal movement, resulting in more volatile predictions of the optimal gait and therefore more variable steps. This can speed learning of new optimal strategies in new contexts, reducing the penalty due to the accumulated cost of suboptimal movements during learning. But, there is also a penalty to this high motor variability—once the new optimal strategy is learned, motor variability around this optimum means most movements are suboptimal. The optimal solution to this trade-off depends on how quickly the context is changing (**Fig 7**). It is better to learn quickly and suffer steady state variability about the new optimum when the context is rapidly changing. But, when the context changes infrequently, it is better to learn slowly and more effectively exploit the cost savings at the new optimum. Interestingly, the learning rate in our models, which we chose to match our experimental constraints, is optimal for a cost landscape that is changing approximately every 10-15 minutes, a rate of change not dissimilar from that applied in our experiment protocol. In humans, the nervous system likely has control over the learning rate and the amount of exploration, and may adjust both based on its confidence in the constancy of the energetic conditions. This suggests exploration, and potentially faster learning, could be promoted not through consistent experience in an energetic context, but rather by experimentally alternating energetic contexts.

**Fig 7.**
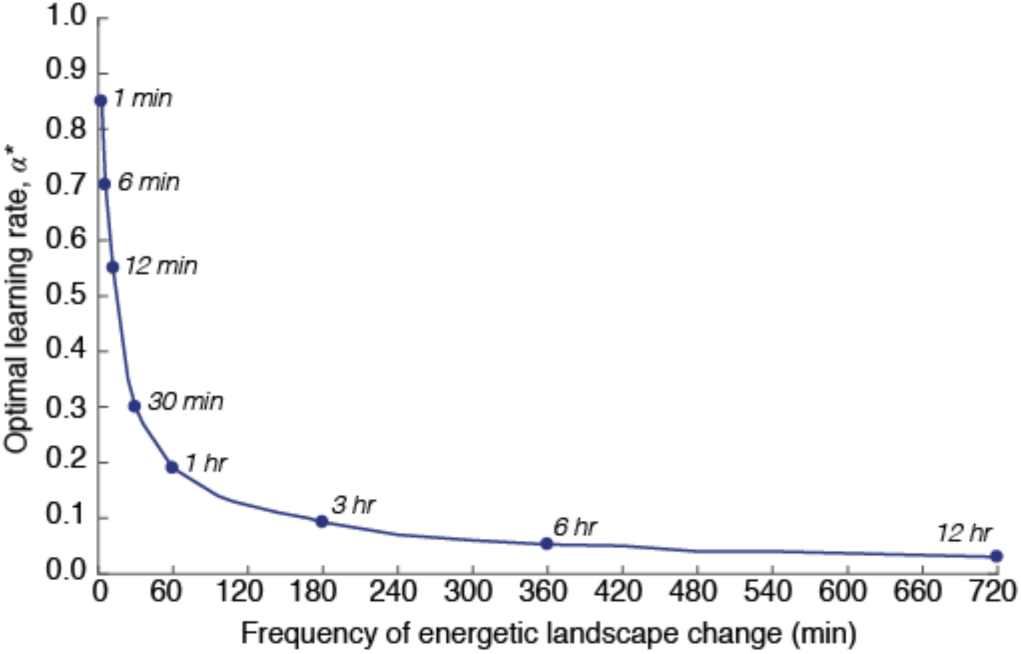
Energetically optimal learning rates for varying frequency of cost landscape change. Measurement and execution noise were set at 2.0% and 1.5%, respectively.

Identifying the dimension of an optimization problem may be the trigger for initiation. The coordination of walking is a task of dauntingly high dimension [17,50]. Various gait parameters, including walking speed, step frequency, and step width must be selected, and numerous combinations of muscle activities can be used to satisfy any one desired gait. When presented with new contexts, the nervous system must identify which parameters, if any, to change in order to lower cost. The difficultly of this task may partly explain why non-spontaneous adaptors do not initiate optimization when the exoskeletons are turned on and they are immediately shifted to a higher cost gait. Although it may be clear to the nervous system that costs are higher, it may remain unclear how it should change movement to lower the cost. This could also explain why in some past experiments, by our group [31] and others [27,51], people did not initiate optimization and discover new energy optimal coordination strategies. Experience with lower step frequencies, and therefore lower costs, may allow the nervous system to identify that this is the relevant dimension along which to optimize. This behavioural phenomenon is captured by the addition of a reference cost to our simple reinforcement learning algorithm, and has parallels in classic feedback control models as well as neurophysiological habituation [13,52,53]. Our experiments have demonstrated how the nervous system rapidly solves a one-dimensional optimization problem—where we alter the energetic consequences of a single gait parameter and apply targeted experience along this dimension of gait. How the identified mechanisms extend to optimizing higher dimension movement problems, like those often encountered in real-world conditions, remains an open question for future work.

Unveiling the mechanisms that underlie the real-time learning of optimal movements may indicate how this process can be accelerated. This has direct applications in the development of rehabilitation programs, the control of assistive robotic devices, and the design of sport training regimes. A stroke patient faced with a change to their body, a soldier adapting to the new environment created by an exoskeleton, and an athlete attempting to learn a novel task, all seek new optimal coordination strategies. Our findings indicate that eliciting exploration through high motor variability, as well as targeted experience along the relevant movement dimension could rapidly accelerate motor learning in these circumstances by cueing the nervous system to initiate optimization. Therapists and coaches may commonly be doing just this, based on years of accumulated knowledge about effective learning strategies. In this view, a more mechanistic understanding of the nervous system’s internal algorithms could aid therapists and coaches in setting a course for a patient or athlete to navigate through various possible movement strategies.

## METHODS

### Subjects

Testing was performed on a total of 36 healthy adults (body mass: 63.9 ± 9.8 kg; height: 1.69 ± 0.10 cm; mean ± SD) with no known gait or cardiopulmonary impairments (Table S1). Simon Fraser University’s Office of Research Ethics approved the protocol, and participants gave their written, informed consent before experimentation.

Initially, 27 subjects were randomly assigned to one of three experiments (9 subjects per experiments) in which their second experience period included either included either metronome-guided experience with discrete cost points on a new cost landscape (**Fig 2D**), metronome-guided experience with many costs along a new cost landscape (**Fig 2E**), or self-guided experience many costs along a new cost landscape (**Fig 2F**). A preliminary analysis revealed that 5 of the 27 subjects (1 from the metronome-guided discrete experience experiment, 1 from the metronome-guided broad experience experiment, and 3 from the self-guided broad experience experiment) appeared to gradually adapt their gait toward the optima during the first experience period, prior to second experience period (**Fig 2C**). These subjects, whom we refer to as ‘spontaneous initiators’, were therefore not included in the analysis for the second experience periods and were instead analyzed as a separate group. To be considered a spontaneous initiator subjects had to meet two criteria. First, during the final 3-minutes of the first experience period their average step frequency (final step frequency) was required to fall below 3 SD in steady state variability, determined from the final 3-minutes of the baseline period. For most subjects, this equates to a minimum decrease in step frequency of approximately 5%. Second, the decrease in step frequency could not be an immediate and sustained mechanical response to the exoskeleton turning on. The final step frequency had to be significantly lower than the step frequency measured in the 10th to 40th second after the exoskeleton turned on (one-tailed t-test, p>0.05). To rebalance experiments, an additional 2 subjects, both of whom were non-spontaneous initiators, were added to the self-guided broad experience experiment.

An analysis of data from the second experience periods indicated that the experience with either high and low step frequencies, and therefore costs, prior to the probe had a lasting effect on the subjects’ self-selected step frequency during the probe. To investigate the interaction between high and low cost experience, as well as the order of the experience, we wanted to compare the time course of adaptation during the first and final probes. To achieve the statistical power necessary to do so, we added an additional 7 subjects to the metronome-guided discrete experience experiment. For the added subjects the experience prior to the first or last probes were set to be either the highest (+10%) or lowest (−15%) step frequency, with all other step frequencies assigned in random order. One of the added subjects met the criteria for a spontaneous initiator and was therefore not included in this investigation between experience direction and time. In total for the analysis, 5 subjects experienced +10% and 4 experienced - 15% prior to the first probe. Prior to the last probe, 4 subjects experienced +10% and 5 experienced −15%. While these subject numbers are low, to detect an across subject average difference in step frequency of at least 5%, given across subject average standard deviation in step frequency of 2.5%, we calculated that we required only 4 subjects per group to achieve a power of 0.8. In addition, we see clear trends in individual subject data (**Fig S2**).

In total, 6 of the 36 tested subjects were identified as spontaneous initiators (body mass: 60.8 ± 10.6 kg; height: 1.68 ± 0.11 cm; mean ± SD), while 14 were included in the analyses for the metronome-guided discrete experience experiment (body mass: 63.0 ± 10.7 kg; height: 1.69 ± 0.11 cm; mean ± SD), 8 for the metronome-guided broad experience experiment (body mass: 67.7 ± 9.1 kg; height: 1.71 ± 0.09 cm; mean ± SD), and 8 for the self-guided broad experience experiment (body mass: 64.2 ± 8.6 kg; height: 1.67 ± 0.07 cm; mean ± SD).

### Detailed Experimental Protocol

Each subject completed one testing session, lasting three hours with no more than two and half hours of walking to reduce fatigue effects. All subjects experienced the penalize-high control function, which has previously been shown to shift energetic optima to low step frequencies ***[31]*** (**Fig 1C-G**). Subjects were given between 5-10 minutes of rest in between each of the walking periods, including baseline, habituation, first experience, and one of the three assigned second experience periods (**Fig 2A-F** respectively, described in detail below).

At the beginning of testing, we instrumented subjects with the exoskeletons and indirect calorimetry equipment (VMax Encore Metabolic Cart, VIASYS®). We then determined their resting energetic cost during a 6-minute quiet standing period. Following this, during the baseline period (**Fig 2A**), subjects were simply instructed to walk for 12-minutes.

Next, subjects completed a habituation period where they were familiarized with walking at a range of step frequencies (**Fig 2B**). This trial was included to reduce the possibility that in future trials subjects could be unaware they are able to alter, or fearful to alter, their step frequency when walking on a treadmill at a constrained speed (**Fig 2B**). During this habituation, the controller remained off; therefore, subjects did not gain experience with the new energetic landscape. We explained to the subjects that for a given walking speed it is possible to walk in a variety of different ways, including with very long slow steps or very short fast steps. They were encouraged to explore these different ways of walking during the habituation period. They were also informed that at times a metronome would play different steady state tempos, or slowly changing tempos, and that they should do their best to match their steps to the tempos. During the habituation period, three different steady state tempos were played for three minutes each. These tempos included +10%, −10%, and −15% of the initial preferred step frequency. The sinusoidally varying metronome tempo had a range of ±15% of the initial preferred step frequency and a period of 3 minutes.

Prior to the first experience period, we explained to subjects that they would next walk for 6 minutes with the exoskeleton turned off, at which point the exoskeleton would turn on and they would walk for a remaining 12 minutes (**Fig 2C**). They were given no other directives about how to walk and at no point during testing were subjects provided with any information about how the controller functioned, or how step frequency influenced the resistance applied to the limb.

For the second experience period, subjects completed one of the three experiments: metronome-guided experience with discrete cost points on a new cost landscape (**Fig 2D**), metronome-guided experience with many costs along a new cost landscape (**Fig 2E**), or self-guided experience many costs along a new cost landscape (**Fig 2F**). All subjects were informed that they would be walking for 30 minutes and that the exoskeleton would be on for the first 24 minutes and off for the final 6 minutes. To gain insight into the progress of optimization under each experiment, 1-minute probes of subjects’ self-selected step frequency occurred at the 6^th^, 10^th^, and 14^th^ minute, along with a final 6-minute probe at the 18^th^ minute (**Fig 2D-F**).

Those assigned to the metronome-guided discrete experience experiment were informed that at times the metronome would be turned on, during which they should match their steps to the steady-state tempo, and that when the metronome turned off, they no longer had to remain at that tempo (**Fig 2D**). Besides these instructions, subjects were given no further directives about how to walk. The metronome was turned off at four different time points, each serving as a probe of subjects self-selected step frequency. Prior to each probe, different metronome tempos were played, including −15%, −10%, +5% and +10% of initial preferred step frequency. We chose these tempos such that they spanned the energetic landscape but did not include step frequencies explicitly at the expected optima or the preferred step frequency (approximately −5% and 0%, respectively). Order of the tempos was randomized. The exception to this was that for the 7 subjects added to this experiment, either the first or last tempo was randomly assigned as either +10% or −15%, with the remaining 3 step frequencies assigned in random order.

Those assigned to the metronome-guided broad experience experiment were given the same instructions as those in the metronome-guided discrete experience experiment, except that they were informed that the metronome tempo would change slowly over time (**Fig 2E**). A sinusoidally varying metronome tempo was played for 3 minutes, 4 separate times, which were once again separated by probes of self-selected step frequency. The sinusoidal tempo had a range of ±15% of the initial preferred step frequency, a period of 3 minutes, and began and therefore ended at 0% of the initial preferred step frequency. These subjects were therefore guided through the complete landscape but always returned to their preferred gait prior to a probe.

Those assigned to the self-guided broad experience experiment were informed that at times the experimenter would verbally give them the command ‘explore’, at which point they should explore walking at a range of different step frequencies (**Fig 2E**). They were informed that they should continue to do so until given the command ‘settle’, at which point that should settle into a steady step frequency. They were given no directives about what their steady state step frequency should be. Subjects were instructed to explore four separate times, each lasting three minutes and once again separated by probes of self-selected step frequency. When the command settle was given subjects could be at any self-selected step frequency, therefore the experience direction, at high or low cost, was not predetermined.

### Experimental Outcome Measures

Each subject’s initial preferred step frequency was calculated as the average step frequency during the final 3 minutes of the baseline period. Individual subject’s variability in step frequency, calculated as a coefficient of variation, was also assessed during this time period. Similarly, the average step frequency was calculated during the final 3 minutes of the first experience period. During this period, the spontaneous initiators were found to adapt toward the optima. To determine the average rate at which they did so, step frequency time series data from the 6^th^ to the 18^th^ minute for the subjects was grouped together and fit with a single term time-delayed exponential. Prior to fitting, data was down-sampled to a step rate of 1.8Hz, so as not to overestimate data points and inflate calculated confidence intervals. We used one-tailed t-tests with a significance level of 0.05 to compare the step frequency, as well as variability in step frequency, of the spontaneous and non-spontaneous initiators (**Fig 3C-D**). We used one-tailed t-tests because we expected the spontaneous initiators to present with lower steady-state step frequencies and higher variability.

During the second experience periods, 1-minute probes of subjects’ self-selected step frequency occurred at the 6^th^, 10^th^, and 14^th^ minute, along with a final 6-minute probe at the 18^th^ minute. When statistical comparisons were made between first and last probes following high and low experience, data from the 30^th^ to the 60^th^ second of each of the self-selected step frequency probes were used for analysis. We used t-tests with a significance level of 0.05 (**Fig 4**, **Fig 5B** and **Fig 5D**). When investigating the rate at which subjects adapted their step frequency, step frequency time series data from the first 60 seconds of the probes from subjects of the same experiment were once again fit with a single term time-delayed exponential, using the same process steps as previously described. For plotting purposes, we averaged across subjects of the same experiment and calculated the across subject standard deviation at each time point.

Because there was no effect of experience direction during the last probe, subjects from the high and low experience were grouped. The final preferred step frequency was calculated as the average step frequency during the 21^st^ to 24^th^ minute of the second experience period, just prior to the controller being turned off. The re-adaptation step frequency was calculated as the average step frequency during the final 3 minutes of the second experience period, when the controller was turned off. When investigating the rate at which subjects re-adapted their step frequency back to the initial preferred, step frequency time series data from the entire re-adaptation period were once again fit with a single term time-delayed exponential and the average and standard deviation profiles were calculated for plotting purposes.

### Model Details

The range of possible actions (*a_i_, a_i+1_, … a_n_*) were set to be discrete integer step frequencies, ranging between −15% and +15%. The simulated energetic cost landscape (*Q\**), before the controller was turned on, was modelled as a quadratic function of the form:

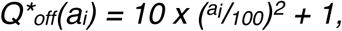

having a normalized cost of 1 at the optimum and a curvature that well approximates our experimentally measured landscape. After the controller was turned on, the simulated landscape was modelled as:

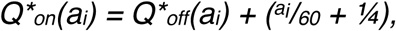

where the cost added to *Q*_off_(a_i_)* well approximates the energetic effect of our controller, again creating a landscape similar in shape to that which we measure experimentally.

The parameters that describe the behavior of the reinforcement learner include: the standard deviation in step frequency execution noise (*n_e_*), the standard deviation in energetic cost measurement noise (*n_m_*), and the weighting parameter, or learning rate (*α*). On any given step, the levels of measurement and execution noise are drawn from a Gaussian distribution with mean zero and standard deviation determined from the value of the parameter.

We performed a sensitivity analysis to determine feasible parameter ranges that are consistent with experimentally measured rates of convergence to the optimum and variability in step frequency. Similar to the design of our experimental first experience period, we simulated a protocol that lasts 1440 steps (approximately 12 minutes) in which the landscape changed from *Q*_off_* to *Q*_on_* after 720 steps (approximately 6 minutes). Using our simple reinforcement leaning model that spontaneously initiates optimization, we varied the execution noise (between 1 and 3% of the initial preferred step frequency), measurement noise (between 0.1 and 6% of the energetic cost at the initial preferred step frequency during natural walking), and learning rate (between 0.01 and 1). Model simulations were repeated 1000 times for each possible combination of parameter settings.

We then determined the resulting rates of convergence to the optimum by averaging step frequency data across repeats and then fitting the final 720 steps with a single process exponential model. As expected, higher learning rates, which put greater weight on new measurements as opposed to past measurements, lead to faster convergence to the optimum (**Fig S3A**). This rate of convergence is largely unaffected by measurement noise, and is only minimally affected by execution noise, where higher execution noise can slow convergence to the optimum. In our experiments, the convergence to the optimum typically occurred with a time constant of less than 100 steps. This experimental constraint leaves us with a wide range of possible learning rate parameter settings, from 0.5-1.0 for any simulated combination of measurement and execution noise. For the purposes of our simulations, we set the learning rate to be 0.5.

Given our chosen learning rate, we next selected measurement and execution noise levels that generated variability in step frequency that well approximated that which we observed experimentally during steady state behavior (1.0-1.5%). For each simulation repeat, we calculated the standard deviation in step frequency during the first 720 steps. During this time, the learner is at the *Q*_off_* optimum and the landscape is unchanging, leading to steady state behavior. We then averaged this value across repeats to get an average measure of variability in steady state step frequency for each possible combination of measurement noise and execution noise. Once again, our experimental constraints left us with a wide range of possible parameter settings (**Fig S3B**). For the purposes of our simulations, we set the measurement noise to be 2.0% and the execution noise to be 1.5%. Importantly, within the ranges deemed reasonable by our experimental constraints, the qualitative behaviours generated by our model are not particularly sensitive to the specific learning rate, measurement noise, and execution noise parameter settings we chose.

Our reference cost model only initiates optimization after sufficient experience with a lower cost gait. It is unclear from our experiments exactly what constitutes sufficient experience with a low-cost gait. For example, it may require a substantially lower cost, a sufficient number of steps at a lower cost, or some combination of these criteria. For the purposes of modelling, we assume that the criteria have been met during the experience with low cost prior to the first probe, in keeping with our experimental findings.

Principles of energetic optimality may also determine the choice of learning rate. It is possible to solve for a learning rate that minimizes energy expenditure; however, the optimal learning rate is dependent on how frequently the energetic landscape is changing. To demonstrate this, we simulated protocols where the landscape changes from *Q*_off_* to *Q*_on_* with a period varying between 1 minute and 12 hours, at a duty cycle of 50%. We simulated 24 hours of walking and evaluated learning rates ranging between (between 0.01 and 1). Model simulations were repeated 100 times for each possible combination of parameter settings. We then determined the average energetic cost across all steps (before measurement noise was applied), and then averaged across repeats to get an average energetic cost for each combination of period and learning rate. Next, we solved for the learning rate that minimized energetic cost for each period (**Fig 7**).

### Alternative Models

Although our reinforcement learning model is quite simple in form, it is reasonable to ask if even simpler algorithms could capture our experimental behaviour. We first considered models that lacked two key features of our final model—the storing of the entire value function and the need to update a reference cost prior to initiation of optimization.

A simplified model that forgoes the storing of the entire value function can reproduce the key features of our experimental data. This simplified no-value function model only stores the optimal gait and its associated cost, rather than the costs at all experienced gaits (i.e. the value function). Yet, it can initiate optimization after experience with a lower cost, converge on a new energetic optimum using a local search, and learn to rapidly predict this optimum when perturbed away. Despite this, we prefer the slightly more complex value-function model because we suspect it will better generalize to learning in the real world. We suspect this for two reasons. First, storing information about non-optimal gaits seems valuable given that at times one may be constrained from using the globally optimal gait. For example, the no-value function model would need to relearn the optimal walking speed when constrained by a slow crowd that prevents walking at the globally optimal speed. In contrast, the value function model, which has memory of past non-optimal walking experience, could rapidly predict the new cost optimal speed in the face of this constraint [54]. Ignoring this potentially useful past experience seems unlikely on the part of the nervous system, given that there will be times when it is energetically beneficial to recall it. Second, the simpler model avoids a value function only in the case where the learning task has one dimension, such as in our experimental paradigm. If instead, for example, the nervous system had to learn the optimal speed and step frequency it would need to store the optimal step frequency, and its cost, at each speed. This is a one-dimensional value function for a two-dimensional optimization problem. As the nervous system can’t know a priori the dimensionality of the optimization problem, it may benefit from learning a high dimensional value function and then constraining the optimization problem depending on the constraints of the task.

A simplified model that forgoes the updating of a reference cost prior to initiation of optimization cannot reproduce key features of our experimental data. In our model of non-spontaneous initiators, prior to initiation of optimization, the learner only updates a reference cost (**Fig 1B**). Without this feature, direct and gradual convergence to the new energetic optimum after forced experience with a low cost is not produced. Instead, because the reference cost has not been updated and therefore is expected to be that experienced under the controller off condition (*Q*_off_*), this model will first rapidly shoot back to the old cost optimum after experience with a low cost. Only after updating this cost estimate, to its now higher cost value under *Q*_on_*, will it then gradually adapt to the new optimum. This updating of a reference cost prior to initiating optimization is not only necessary to reproduce our experimental findings, but also has many parallels in neurophysiological habituation [13,52,53].

In all our models, we chose to discretize the possible actions, or step frequencies. This enforces local learning, where actions at distinct step frequencies have no effect on the expected value of others. It is entirely possible, if not likely, that the nervous system does not discretize its action space in this way but may rather store a function. In other words, the value function may not be a lookup table, as we have modeled it, but rather some representative equation, such as a polynomial. However, it still seems likely that the nervous system updates its value function in some localized way, as global function approximations are known to produce highly variable behavior away from the local area of learning [55].

## ACKNOWLEDGMENTS

This work was supported by a Vanier Canadian Graduate Scholarship (J.C.S.), a Michael Smith Foundation for Health Research Fellowship (J.D.W.), the U.S. Army Research Office grant #W911NF-13-1-0268 (J.M.D.), and an NSERC Discovery Grant (J.M.D.). We thank T.J. Carroll and R.T. Roemmich for their helpful comments and suggestions.

## AUTHOR CONTRIBUTIONS

J.C.S. and J.M.D. designed the study with input from J.D.W. and S.N.S.; J.C.S. collected data and performed analysis; J.C.S. and J.M.D. wrote the manuscript. All authors discussed the results and commented on the manuscript.

## SUPPORTING INFORMATION CAPTIONS

**Table S1.**
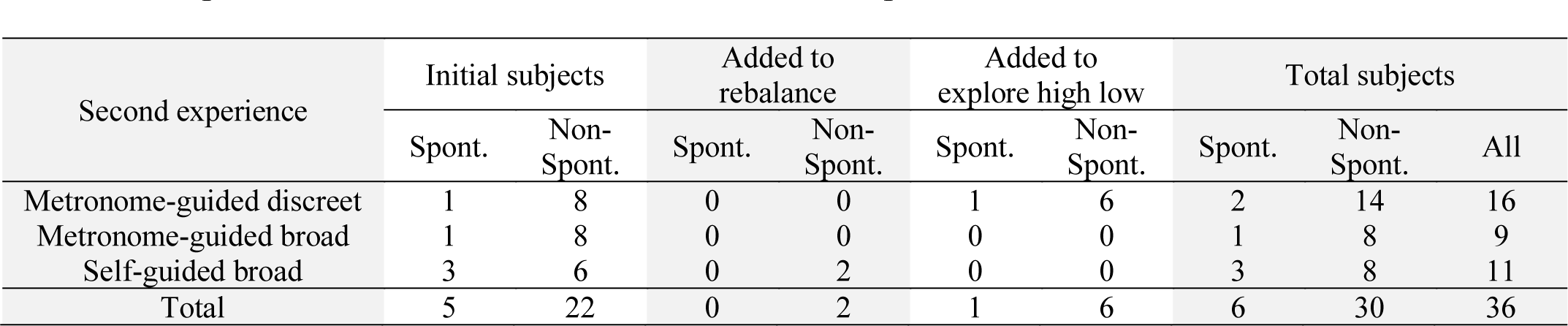
Subject numbers per experiment. We initially tested 9 subjects in each of the three Second Experience testing periods. To account for a high number of spontaneous initiators in the self-guided broad experience condition we added an additional 2 subjects to this group to rebalance our conditions. To achieve the statistical power necessary to investigate the interaction between high and low cost experience, as well as the order of the experience, we added an additional 7 subjects to the metronome-guided discrete experience experiment, one of which we found to be a non-spontaneous initiator. In total, we tested 36 subjects, 6 of which were classified as spontaneous initiators and 30 which were non-spontaneous initiators.

**Fig S1.**
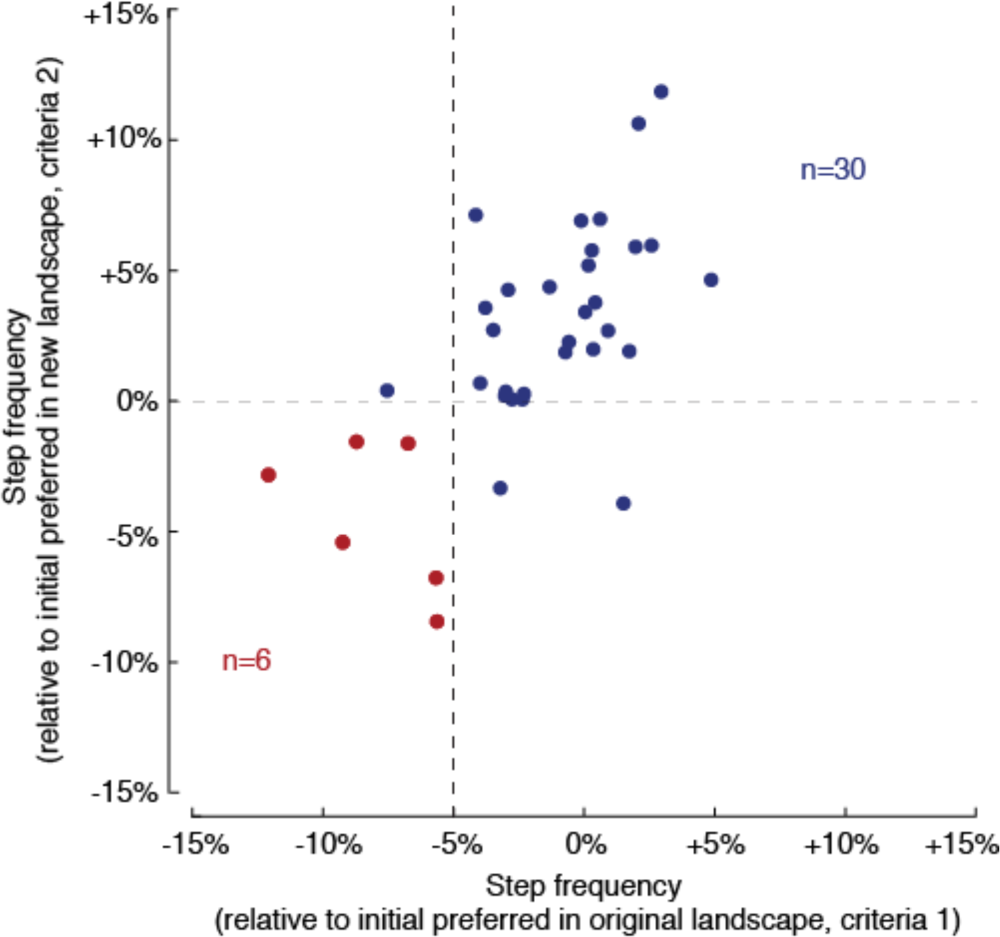
Discrimination plot of spontaneous and non-spontaneous initiators. We defined spontaneous initiators as having a first experience final step frequency consistent with the expected optima (−3SD from the initial preferred step frequency, or approximately −5%, x-axis), as well as displaying a significant change in step frequency from that displayed immediately after the exoskeleton was turned on (significantly different from 0%, y label). Although the above statistics, and not simple thresholds, were used for each criteria, the dashed lines illustrate roughly how each criteria divided the data.

**Fig S2.**
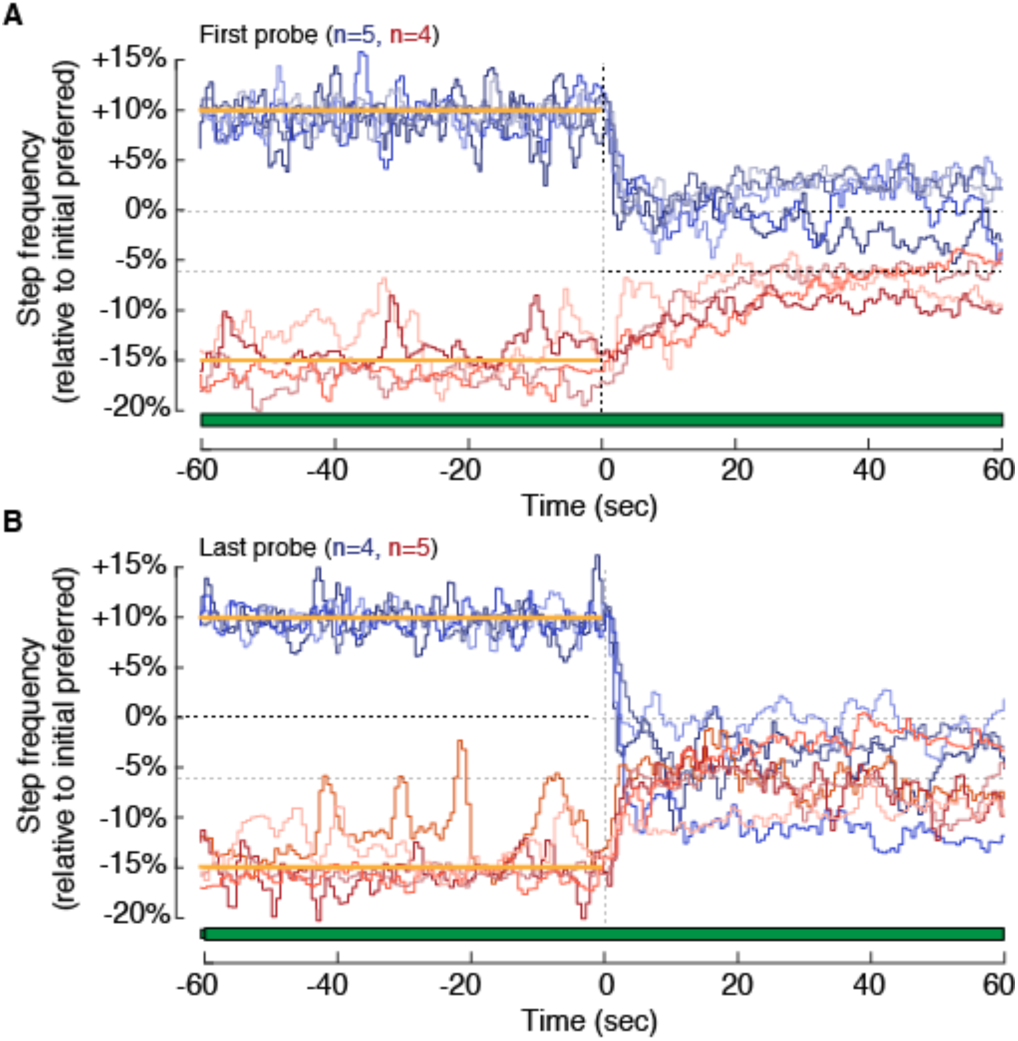
Individual subject effect of experience direction during first and last step frequency probe. Step frequency time-series data for the first (**A**) and last (**B**) probes following experience either high (blue) or low (red) step frequencies for individual subjects. The horizontal bars indicate when the controller is turned on (green fill) and off (white fill), and the yellow lines indicate the prescribed metronome frequencies.

**Fig S3.**
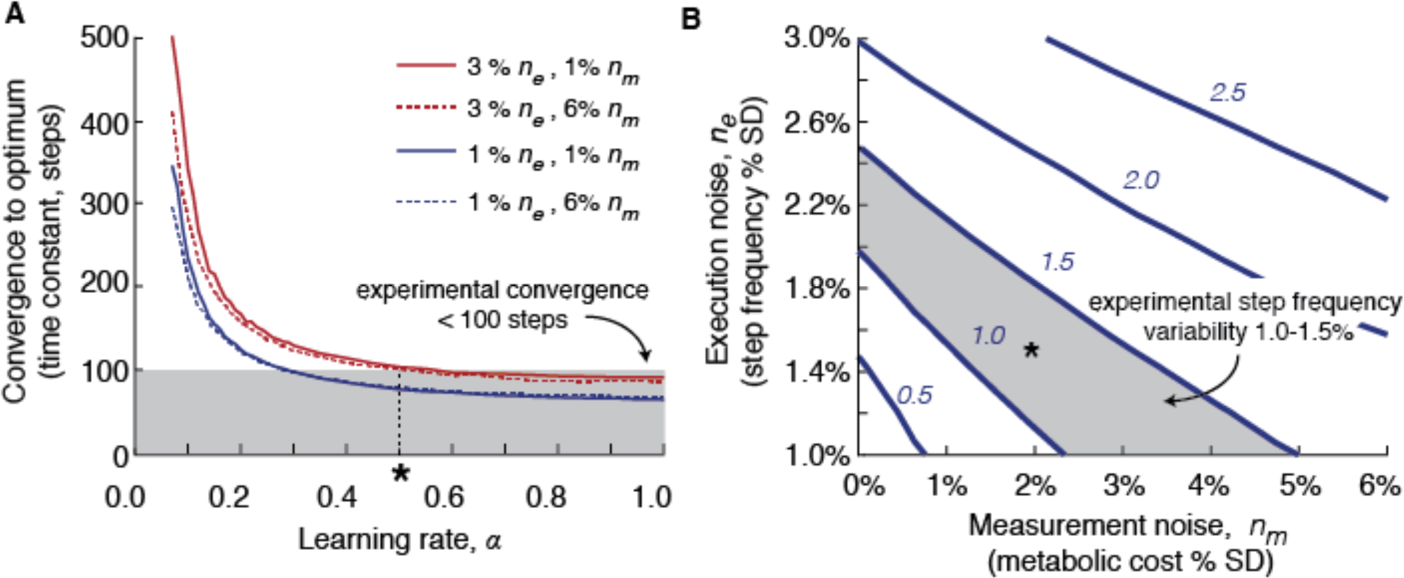
Sensitivity analysis of model parameters. (**A**) Effect of varying the learning rate parameter on the rate of converge to the energetic optimum for different measurement and execution noise levels. The shaded region represents a reasonable convergence rate given that observed experimentally (maximum 100 steps), while the asterisk and dashed vertical line represents the chosen learning rate parameter value used in simulation (0.5). (**B**) Effect of varying measurement and execution noise on variability in steady state step frequency. Learning rate was kept constant at 0.5. Each line and the associated italic number represents a constant value of steady state step frequency. The shaded region represents reasonable steady state step frequencies given that observed experimentally (1.0%-1.5%). The asterisk represents the chosen measurement and execution noise parameter values used in simulation (2.0% and 1.5%, respectively).

